# Interferon Beta Drives Therapy Resistance in a Patient-Derived Model of High-Grade Serous Ovarian Cancer

**DOI:** 10.64898/2026.01.27.699126

**Authors:** Ashlyn Conant, Tise Suzuki, Kiera McGivney, V S S Abhinav Ayyadevara, Sharon Asariah, Jay Deng, Ethan Nyein, Jacqueline Coats, Gary Yu, Yevgeniya Ioffe, Christian Hurtz, Juli J. Unternaehrer

**Author notes:** **Corresponding Author:** Juli J. Unternaehre, Associate Professor, Department of Basic Sciences, Division of Biochemistry, Loma Linda University School of Medicine, 11085 Campus Street, Mortensen Hall 219, Loma Linda, CA 92354.

## Abstract

Cancer cell-autonomous type 1 interferon (IFN-1) production and signaling is frequently activated in response to DNA damage and has been associated with the development of therapy resistance in several cancer types. However, its cell-autonomous role in driving resistance in high-grade serous ovarian cancer (HGSOC), a disease defined by near-universal exposure to genotoxic therapy as frontline treatment, remains unclear. Specifically, whether IFN-1 functions in HGSOC as only a response to genotoxic stress or can independently act in driving resistance phenotypes has not been studied. Utilizing a syngeneic patient-derived model of cisplatin-sensitive (SE) and -resistant (CR) HGSOC, we demonstrate that chronic cisplatin exposure is associated with enrichment of IFN-1 signaling and the interferon-related DNA damage resistance signature (IRDS). Acute cisplatin treatment elicited dynamic, temporal IFN-1 signaling and responses in both sensitive and resistant cells, indicating a conserved stress response in resistant cells. Chronic, low-level exposure to exogenous IFNβ, in the absence of a DNA-damaging agent, was sufficient to phenocopy several features of chronic cisplatin driven resistance, including reduced therapeutic sensitivity, cell cycle arrest, and decreased proliferation. Notably, IFNβ driven resistance occurred without sustained IRDS or canonical interferon stimulated gene (ISG) induction, revealing alternative mechanisms for IFN-1 mediated therapy resistance. Together, these findings identify IFNβ as a functional driver of the development of resistance-associated phenotypes and highlight cell-autonomous IFN-1 signaling as a potential biomarker for resistance and a therapeutic target in platinum-resistant disease.

## INTRODUCTION

High grade serous ovarian carcinoma (HGSOC) remains one of the deadliest gynecologic malignancies despite a more than 80% response rate to frontline therapy^1,2^. Standard of care consists of surgical debulking followed by 6 cycles of platinum/taxane adjuvant chemotherapy^3^. For patients who are not primary debulking candidates, neoadjuvant chemotherapy, followed by interval debulking surgery (IDS) and adjuvant therapy are utilized. Availability of targeted therapies has been steadily increasing but remains limited; vascular endothelial growth factor (VEGF) inhibitors, such as bevacizumab, and poly-ADP ribose polymerase (PARP) inhibitors, such as olaparib, and antibody drug conjugates (ADCs) such as mirvetuximab being some of the most common therapies for women with specific indications^3-5^. Regardless of the robust frontline response, more than 70% of women experience recurrence and associated therapy resistance, contributing to a bleak >30% 5-year survival rate for stage IV disease^6^. The initial responsiveness coupled with subsequent treatment-resistant recurrence combine to demonstrate the urgent need for reliable models of acquired therapy resistance.

Within tumor cells, genomic instability, DNA damage, and aberrant mitotic events all contribute to the accumulation of cytosolic DNA, which activates the cGAS-STING (cyclic GMP-AMP synthase - stimulator of interferon genes) pathway and promotes type 1 interferon (IFN-1) production^7-10^. This mechanism is also amplified by DNA-damaging therapies, such as chemotherapy, and by the presence of damage-associated molecular patterns (DAMPs) in the tumor microenvironment (TME)^11,12^

The intensity and duration of the IFN-1 signal greatly influence the cellular response and phenotypic adaptation via interferon-stimulated gene (ISG) induction^13^. Canonical ISG expression is driven by a robust and acute IFN-1 signaling event which is transduced via canonical JAK/STAT signaling, leading to the formation and phosphorylation of the interferon-stimulated gene factor 3 (ISGF3) transcription factor complex. This signaling correlates with anti-proliferative, pro-inflammatory, pro-apoptotic, cytotoxic phenotypes^14,15^. Non-canonical ISG signaling, driven by sustained STING activation and chronic low-level IFN-1 production, drives non-canonical JAK/STAT signaling and formation of the unphosphorylated ISGF3 (u-ISGF3) complex. This non-canonical signaling promotes a pro-tumorigenic, immunosuppressive, and anti-apoptotic phenotype that results in cytoprotection^16^. A subset of ISGs termed the interferon-related DNA damage resistance signature (IRDS) is highly associated with resistance to DNA damaging therapies^17,18^. This gene set is a pathologically sustained response to non-canonical IFN-1 signaling and has been found to be present, and potentially predictive, in ovarian cancer as early as detection of a p53 mutation in the fallopian tube^19,20^. However, it remains unclear whether IFN-1 signaling in HGSOC is merely a consequence of DNA damage or whether it can independently drive therapy resistance and phenotypic adaptation.

In this study, we aimed to define the role of cancer cell autonomous IFN-1 production and signaling in the development of therapy resistance, and associated phenotypes, within HGSOC. Using a syngeneic patient derived model of cisplatin-sensitive and -resistant HSGOC, we demonstrate that chronic cisplatin exposure is associated with enrichment of IFN-1 and IRDS signatures. Acute cisplatin treatment is shown to differentially and temporally enhance IFN-1 production, as well as related signaling, in sensitive and resistant cells, highlighting the effect of prior therapy on stress response. In validation of the role of DNA damage-induced IFN-1 activation, sensitive and resistant cells treated with chronic, low-level IFNβ phenocopied key features of cisplatin resistance, including altered cell states and reduced cisplatin sensitivity. Notably, IFNβ driven resistance occurred in the absence of sustained IRDS or canonical ISG induction, indicating the existence of alternative mechanisms by which IFN-1 promotes therapeutic response adaptation. These findings identify IFNβ as a functional driver of cisplatin resistance and associated phenotypes, highlighting cell-autonomous IFN-1 signaling as a potential biomarker and therapeutic target.

## MATERIALS AND METHODS

### Cell Culture

The collection and use of the cells in this study were approved by the Loma Linda University (LLU) Institutional Review Board (IRB, 58328). With informed consent, deidentified tumor tissue was collected by the Loma Linda University Cancer Center Biospecimen Laboratory (LLUCCBL), and immediately transported to the laboratory for processing. PDX samples were collected, preserved, and processed as previously described^21^. The same tumor samples were used to clinically diagnose HGSOC.

Patient derived cells (PDX4 SE/CR) were cultured in a 3:1 mixture of HycloneTM Ham’s Nutrient Mixture F12 with L-glutamine (SH30026.01; Cytiva) and Dulbecco’s Modified Eagle’s Medium with high glucose and L-glutamine (DMEM; 25-501), supplemented with 5% FBS (Omega Scientific), 0.4 μg/mL hydrocortisone (H0888-1G; Sigma-Aldrich), 5 μg/mL insulin (91077C-100MG; Sigma-Aldrich), 2 μg/mL isoprenaline hydrochloride (I5627-5G; Sigma-Aldrich), 24 μg/mL adenine (A8626; Sigma-Aldrich), 100 U penicillin, and 100 μg/mL streptomycin (25-512; Genesee Scientific).

Cells were treated with their respective IC50’s of cis-Diammineplatinum(II) Dichloride (cisplatin; D3371-100MG; TCI Chemicals) based on cell viability assays^21,22^.

Cells were treated with Human Interferon Beta 1a at 100U/mL or 2U/mL (11415-1; PBL Assay Science) diluted in 0.1% BSA in sterile 1x PBS.

### Library Preparation and RNA Sequencing

Total RNA was extracted from PDX4 SE and PDX4 CR cells using the miRNeasy Mini Kit (217004, Qiagen, Germantown, MD, USA). Subsequently, RNA-seq library construction and generation of raw data was performed at the Loma Linda University Center for Genomics using the Ovation Universal RNA-seq System (0364; Tecan; Männedorf, Switzerland). Briefly, 100 ng of total RNA was reverse transcribed and then made into double-stranded cDNA by adding a DNA polymerase. cDNA was concentrated using Agencourt beads, followed by end repair and adaptor ligation. Unique barcodes were used for each sample for multiplexing. Targeted rRNA-depletion was performed before the final library construction. Libraries were amplified using 13 cycles in the Eppendorf™ Mastercycler™ pro PCR system (Hamburg, Germany) and purified using Agencourt beads. RNA-seq libraries were sequenced on Illumina NextSeq 550 (Illumina, San Diego, CA, USA) with single 76 bp reads. Illumina RTA v 2.4.11 software was used for basecalling, and bcl2fastq v 2.17.14 was used for generating FASTQ files.

Specific analysis pipelines and codes used to generate graphs can be found at https://github.com/tsuzukiPhD/RNAseq (accessed on 8 January 2024), and bioinformatics analysis was done as described ^22^.

### Pathway Analysis

Ingenuity Pathway Analysis (IPA; QIAGEN Inc., Hilden, Germany, https://digitalinsights.qiagen.com/IPA; accessed on 5 May 2023) was used for the identification of canonical/hallmark pathways activated among the DEGs obtained through the PDX4 RNA-seq analysis^23^. To identify specific activated pathways associated with interferon signaling and JAK/STAT, the following keywords were used for filtering: “interferon” and “JAK”. Bar charts were created with GraphPad Prism v 10.2.2.

Gene Set Enrichment Analysis (GSEA; v 4.3.2; accessed on 27 October 2023) was also used for the identification of pathways activated in the DEGs of PDX4 CR^24,25^. Both Hallmark Gene Sets and Gene Ontology (GO): Biological Processes pathways were queried. Since gene set permutation was used, only pathways with an FDR of less than 0.05 were considered for further analysis. Permutations for GO were set at 1000, and Hallmark Gene Sets at 50,000. Dot plots were created with ClusterProfiler^26^ and display the results of a Gene Ontology (GO) enrichment analysis for the “activated” and “suppressed” DEGs.

### Quantitative Real Time Polymerase Chain Reaction (RT-qPCR)

RNA from cells was isolated using the IBI Isolate DNA/RNA Reagent Kit (IB47602, IBI Scientific, Dubuque, IA, USA) according to the manufacturer’s recommendations. cDNA was synthesized from either 500 ng or 1 μg of total isolated RNA using Maxima First Strand cDNA Synthesis Kit (K1672; Thermo Fisher Scientific). RT-qPCR was performed using Applied Biosystems™ PowerUP™ SYBR™ Green Master Mix (A25778; Thermo Fisher Scientific) and gene primers (Sup Methods, Table 1; custom ordered as cDNA oligos from IDT) on a Stratagene Mx3005P Instrument (Agilent Technology, Santa Clara, CA, USA). The results were analyzed using the Δ cycles to threshold (ΔCt) method.

### IFN-α/β Protein Detection

Secreted IFNα/β was detected using an interferon stimulated gene factor 3 (ISGF3) - secreted alkaline phosphatase (SEAP) HEK-Blue IFNα/β reporter cell line (hkb-ifnabv2; InvivoGen). Concentrations of IFNα/β were calculated by measuring absorbance at 620 nm on a Biotek Synergy H1 Multimode plate reader (Agilent Technology, Santa Clara, CA, USA). A standard curve was generated using a two-fold serial dilution of IFN-β starting at 50 U/mL and plotted with hyperbolic non-linear regression on GraphPad Prism v10.5.0.

### Cell Viability Assay

Cell viability assays were completed using thiazolyl blue tetrazolium bromide (MTT; 00697; Chem-Impex) assays as previously described^21,22^.

### Flow Cytometry

Permeabilized flow cytometry was completed as follows. Cells were stained with BD Horizon™ fixable viability stain 780 (565388; BD) according to the manufacturer’s recommendations for compensation and live cell gating. A single heat shocked sample was used as a positive gating control for FVS. Cells were then permeabilized using 90% ice-cold methanol for 1 hour. Samples were brought to room temperature before rinsing 3x with FACS stain [1% FBS, 0.1% sodium azide (NaN_3_; s2002-5g; Sigma), and 2 mM EDTA in PBS]. Cells were labeled using two staining methods. Cells were stained with directly conjugated fluorescent dye antibodies against STAT1 AlexaFluor™ 647 (558560; BD) and pSTAT1 PE (612564; BD), incubated at room temperature for 30 min, then washed and resuspended in FACS stain. Cells were also stained using a two-step approach first staining with IFITM1 (99969S; Cell Signaling Technology) diluted in 0.5% BSA for 1 hour at room temperature before rinsing and staining with AlexaFluor™ 647 goat anti-rabbit IgG (H+L) secondary antibody (A21245; Fisher Scientific) at room temperature for 30 minutes, washed, and resuspended in FACS stain. UltraComp eBeads (01-2222; Thermo Fisher Scientific), secondary only, and fluorescence minus one (FMO) stained samples were used for positive and negative gating controls where appropriate.

All flow cytometry was performed on MACSQuant Analyzer 10 (Miltenyi Biotec, Bergisch Gladbach, Germany), and data analysis was performed using FlowJo v10.10.0 (FlowJo LLC, Ashland, OR, USA).

### Cell Cycle Analysis

PDX4 cells were treated with 10ng/μL IFN-β for 72 hours in 6-well plates, followed by IFN-β withdrawal by replacing the spent media with 2mL of fresh culture media. Twenty-four hours later, cells were treated with 10μM BrdU (559619; BD Biosciences) for 30 minutes, 4 hours, or 18 hours. Cells were then harvested using 0.05%Trypsin-EDTA (25200-056, Gibco), and samples were prepared for flow cytometric analysis following the manufacturer’s recommended protocol.

### Microscopy

Cell images were obtained using a Nikon Eclipse Ti microscope (Nikon Instruments, Melville, NY, USA) and μManager v1.4.22 software. Aspect ratio was calculated by measuring and dividing the width and length of 20 individual cells per/photo using ImageJ software (National Institutes of Health, Bethesda, MD, USA).

### Proliferation Assay

Cells were plated at 50,000 cells per well in a 12-well plate. Individual wells were trypsinized at various timepoints up to 120 hours and counted using a trypan blue exclusion cell counting method.

### Statistical Analysis

For all experiments, samples in the same treatment group were harvested from at least three biological replicates and tested in three technical replicates. All values in the figures and text are the means ± SD. Graphs were generated, and statistical analyses were performed using Prism v 10.5.0. Statistically significant differences were determined by unpaired t-tests or paired t-tests, as appropriate, unless otherwise noted. p-values of less than 0.05 were considered significant. Outliers were not removed.

## RESULTS

### Chronic cisplatin exposure induced an IFN-1 associated resistant state

In a previous study utilizing a syngeneic chemo-sensitive and resistant patient-derived model of HGSOC, RNA sequencing revealed JAK/STAT among the most upregulated signaling pathways in a chemoresistant cell line (PDX4 CR) when compared to its sensitive counterpart (PDX4 SE)^22^. Gene ontology (GO) enrichment analysis of differentially regulated biological processes in CR compared to SE samples revealed significant activation of pathways involved in IFN-1 production and signaling (Fig 1A). Additionally, Gene Set Enrichment Analysis (GSEA) and Ingenuity Pathway Analysis (IPA) of the sequencing further indicated that IFN-1 production/response gene sets, along with the JAK/STAT pathway were significantly enriched among the DEGs of PDX4 CR (Fig 1B-D). Analysis of individual sample replicates revealed that cisplatin-resistant samples exhibited marked activation of specific genes associated with IFN-1 production (Fig 1C).

**Figure 1:**
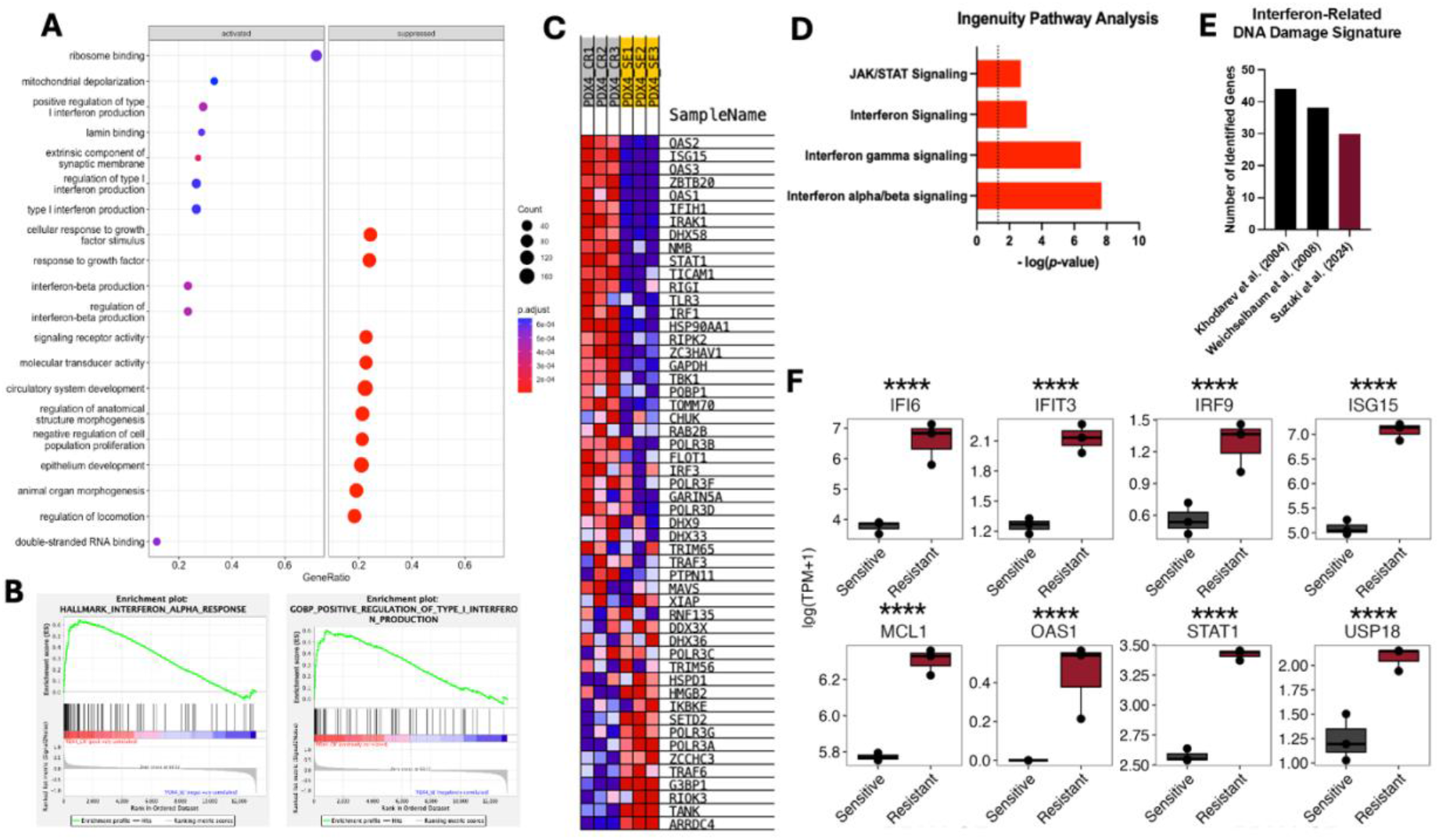
RNA Sequencing of cisplatin sensitive and resistant cells reveals IFN-1 production and signaling signatures. **A.** Gene ontology enrichment analysis of biologic processes activated and suppressed in PDX4 CR compared to SE. **B**. Gene Set Enrichment Analysis (GSEA) of PDX4 cells reveals positive enrichment of the Hallmark Interferon Alpha Response and GO: BP - Positive Regulation of Type 1 Interferon Production Pathways in PDX4 CR. **C**. A GSEA Blue-Pink O’ Gram in the Space of the Analyzed GeneSet histogram displaying the specific genes enriched within the “GO: BP – Positive Regulation of Type I Interferon Production” pathway for the PDX4 samples. The gene expression values are represented in a range of colors, from dark red (upregulation) to dark blue (downregulation). The darker colors correspond to higher differential expression, while the lighter colors correspond to lower differential expression. **D**. Ingenuity pathway analysis (IPA) displaying upregulation of IFN-1 related signaling pathways. **E**. Comparison of the actual number of genes characterized as being involved in the interferon related DNA damage signature (IRDS) expressed in PDX4 CR versus those detailed in Khodarev et al. (2004) and Weichselbaum et al. (2008). **F**. Gene expression profiles of IRDS genes in PDX4 SE (sensitive) and CR (resistant) (Suzuki et al., 2024). p-values: p ≤ 0.0001 (****).

The Interferon Related DNA Damage Signature (IRDS) is a group of genes associated with chemo- and radio-therapy resistance in several cancer cell types^17,18.^ Approximately 30 genes in the IRDS set were identified, compared to those reported in foundational IRDS publications^17,18^ (Fig 1E). Several specific IRDS genes, such as IFI6, IFIT3, IRF9, ISG15, MCL1, OAS1, STAT1, and USP18, were significantly upregulated within the CR cells (Fig 1F). These findings indicate a possible role for IFN-1 signaling in the development of HGSOC chemoresistance.

### IFN-1 production and signaling are transcriptionally engaged, while cytokine output is limited

Candidate genes, including IFN-1 and those associated with IFN-1 signaling pathways, were selected for validation. Total IFNα and IFNβ RNA were significantly higher in PDX4 CR when compared to SE (Fig 2A). Despite the well-known challenges of detecting of IFN-1^27^ we chose to measure the cytokine on the protein level with a bioactive reporter system^28^ which is designed to monitor IFNα/β activity via JAK/STAT activation, as measured by detection of secreted embryonic alkaline phosphatase (SEAP) under the control of ISGF3 binding. In the unstimulated state, without the presence of a DNA-damaging agent, secreted protein expression of IFN-1 protein was similar between SE and CR (Fig 2B). Notably, the levels of IFN-1 protein were close to the lower limits of detection in this assay, suggesting the capacity of cancer cells to produce IFN-1 may be inherently restricted under basal, non-stressed conditions.

**Figure 2:**
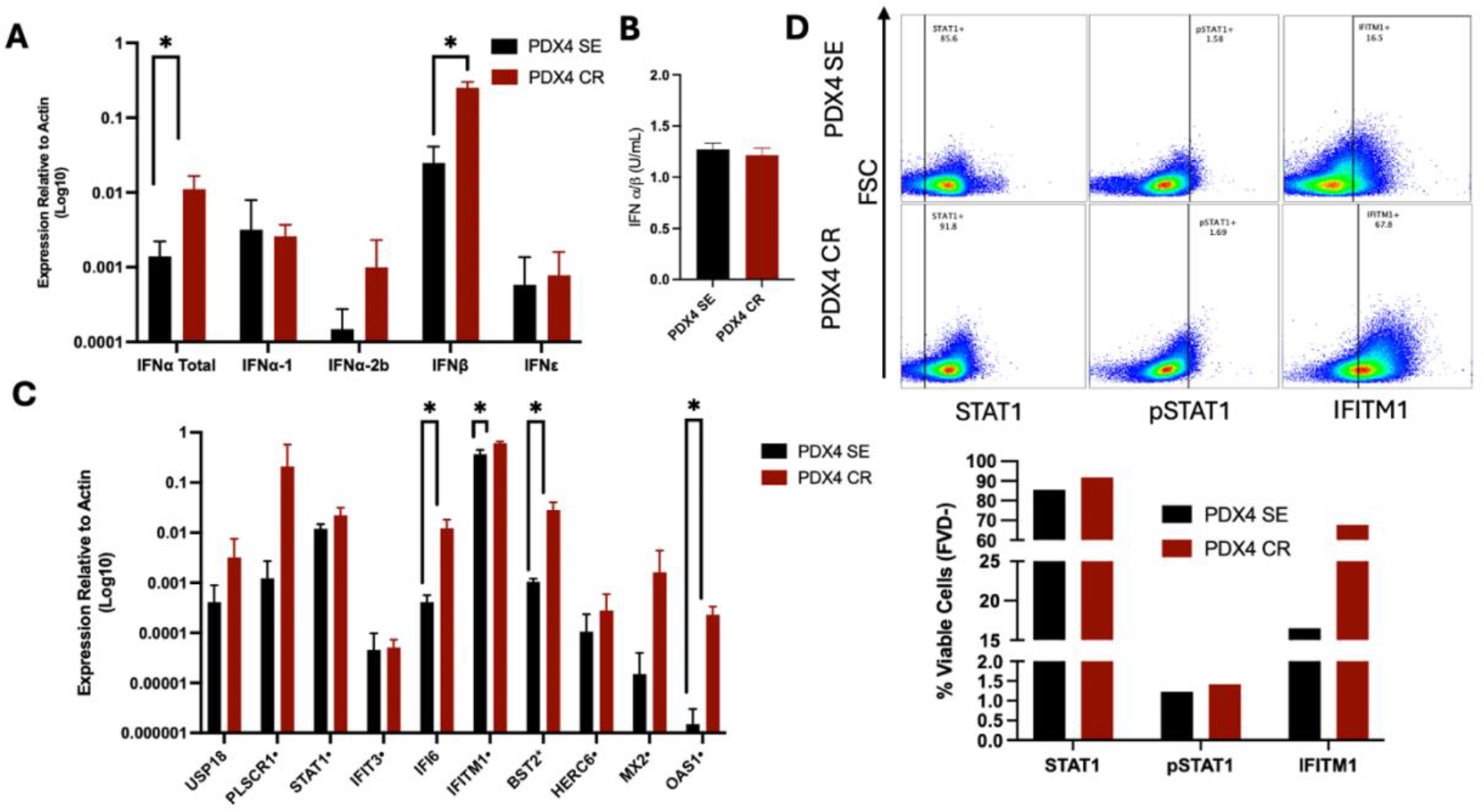
The IFN-I pathway is activated in CR cells. **A.** RT-qPCR analysis of IFN-1 RNA levels. **B**. Secreted IFN-1 protein levels detected in culture supernatant via HEK-Blue SEAP reporter assay. **C**. RT-qPCR analysis of IRDS genes. Dots indicate genes known to be transcribed by u-ISGF3. **D**. Intracellular flow cytometry of IFITM1, total STAT1, and pSTAT1(y701). One representative replicate shown. Results are displayed as n=3, unless otherwise noted, and presented as the means ± SD. Statistical significance was determined using the unpaired t-test. p-values: p ≤ 0.05 (*).

A panel of 10 IRDS genes was selected to further validate the sequencing results. IFI6, IFITM1, BST2, and OAS1 were found to be significantly higher in PDX4 CR compared to SE (Fig 2C). IFI6 (Interferon alpha-inducible protein 6) is implicated in the prevention of apoptosis (via blocking the release of cytochrome C from the mitochondria), mitochondrial regulation (via stabilization of the mitochondrial membrane potential), and DNA replication^29,30^. IFITM1 (Interferon-Induced Transmembrane Protein 1) blocks viral entry into cells (via prevention of viral fusion), inhibits cell growth, and has been found to be required for migration, invasion, and metastasis of several cancer cell types^31-33^. BST2 (Bone Marrow Stromal Antigen 2) plays a potent antiviral role by tethering the lipid envelope of viruses to a host cell membrane and inhibiting viral release and additionally contributes to cell migration^34,35^. OAS1 (2’-5’-oligoadenylate synthetase 1) binds double-stranded RNA (typically from pathogens), activates the production of proteins that stimulate RNases to degrade viral and host RNA, and binds and stabilizes ISGs (e.g. IRF1) and AU rich elements in RNA (e.g. those in IFNβ) to stabilize and prolong RNA lifespan and IFN response^36,37^.

Permeabilized flow cytometric analysis of the cells revealed upregulation of IFITM1 and STAT1 protein in CR cells. pSTAT1, measured as a readout for JAK/STAT signaling activation, was detected with no notable changes in expression between the SE and CR (Fig 2D, gating scheme Sup Fig 1). These data suggest that low-level, sustained IFN-1 signaling, rather than robust cytokine production, may contribute to IRDS enrichment and therapy resistance.

### Acute cisplatin exposure elicits conserved IFN-1 signaling with distinct temporal dynamics in cisplatin-sensitive and-resistant cells

To determine whether sensitive and resistant cells differ in their IFN-1 responses to genotoxic stress, cells were treated with their respective IC50 of cisplatin for 72 hours. Acute cisplatin treatment induced IFN-1 signaling in both PDX4 SE and CR, as evidenced by an increase in secretion of IFNα/β protein (Figure 3A). SE cells displayed an acute IFN response, wherein IFN-1 levels increased at 6 hours. However, CR cells displayed a delayed, sustained IFN-1 response, with IFN levels increasing significantly, compared to SE, in the late phase of treatment. Flow cytometric analysis after 48 hours of cisplatin treatment revealed near universal increases in JAK/STAT signaling activation, as well as IRDS member IFITM1 protein expression regardless of sensitivity or resistance status (Fig 3B).

**Figure 3:**
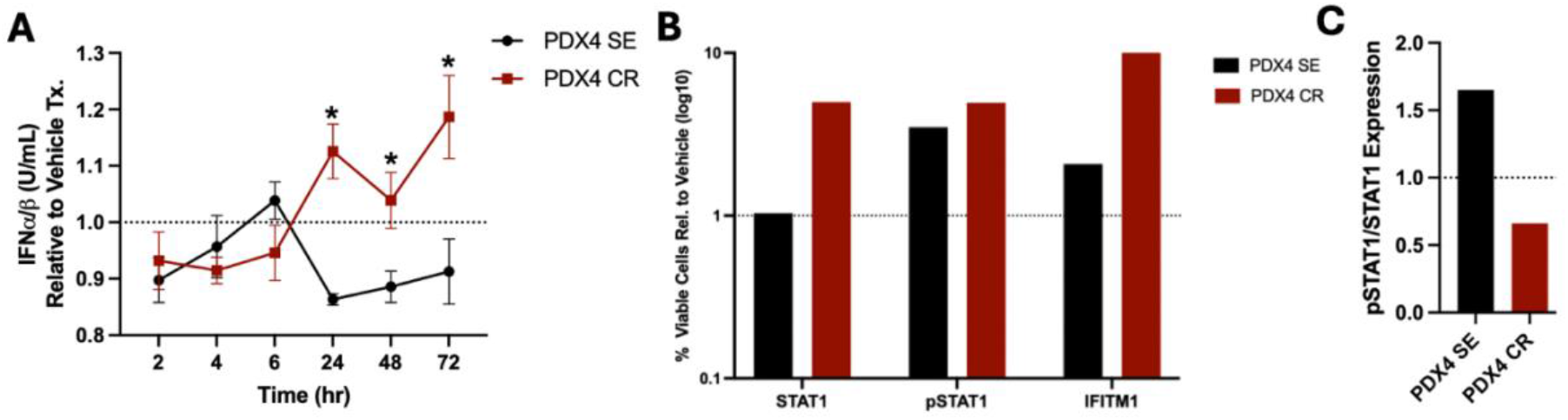
Cisplatin Resistant and Sensitive Cells Respond Differently to Acute Cisplatin Treatment. **A.** Secreted IFN-1 protein levels detected in supernatant, via HEK-Blue reporter assay, collected at various timepoints over 72 hours. Asterisks indicate un-paired T-test comparison between SE and CR samples at each time point indicated. **B**. Flow cytometric analysis of protein expression following 48 hours of IC50 cisplatin treatment. Data are normalized to untreated control cells. One representative replicate shown. **C**. Averaged pSTAT1/STAT1 values from three replicates, normalized to untreated control cells. Results are displayed as n=3, unless otherwise noted, and presented as the means ± SD. Statistical significance was determined using the unpaired t-test. p-values: p ≤ 0.05 (*).

Canonical IFN-1 driven JAK/STAT signaling relies upon phosphorylation and dimerization of STAT1 molecules to form the functional ISGF3 complex which transcribes genes in response to IFN-1 signaling. However, unphosphorylated ISFG3 (u-ISGF3) signaling is a non-canonical signaling pathway that has been found to specifically transcribe many ISGs within the IRDS gene set^20,38^. The ratio of pSTAT1 to STAT1 protein expression was calculated and was used to indicate JAK1 kinase activity and canonical/non-canonical signaling. Following acute cisplatin treatment, there was an observable decrease in kinase activity in CR cells, while there was an increase in SE cells, suggesting differential signaling response (Fig 3C).

RNA was also collected at various time points following treatment. IFNβ expression was significantly decreased, when PDX4 CR was treated for 72 hours, as compared to vehicle treated cells. Alternatively, IFNα expression increased significantly at 24 hours in both CR and SE cells (Sup Fig 2A). IFITM1 and STAT1 were also measured and revealed minimal changes in expression upon cisplatin treatment (Sup Fig 2B).

These data indicate that an IFN-1 response may be a conserved component of cellular response to acute DNA-damage^39^ in HGSOC, while prior exposure to cisplatin may alter the temporal dynamics of this response rather than its overall capacity of activation.

### Chronic, low-level IFN-1 treatment phenocopies chronic cisplatin treatment

As cisplatin resistance has been associated here with sustained IFN-1 signaling and altered temporal responses to acute genotoxic stress, we sought to directly investigate the role of IFN-1 signaling in the acquisition of resistance. To determine if IFN-1 is sufficient to drive phenotypic changes associated with therapy resistance, cells were subjected to chronic, low-level IFNβ treatment over time. High-level IFN treatment can drive anti-viral/cancer effects in cells, while low-level IFN is capable of driving the emergence of pro-tumor phenotypes^16^. The low-dose IFNβ regimen used to model the effects of chronic cisplatin treatment was selected based on measured levels of IFN secretion under basal, unstimulated conditions and following cisplatin treatment (Fig 2B, Fig 3A).

Cells were treated with low-level, 2 U/mL IFNβ every 48-72 hours, before harvest at 12 weeks. Chronic IFNβ treatment resulted in phenotypic changes that closely mirrored those observed following prolonged cisplatin exposure. An MTT cell viability assay revealed IFNβ treated cells exhibited increased resistance to cisplatin (Fig 4A). Both SE and CR cells demonstrated notable morphologic changes following treatment, exhibiting a more epithelial-like shape (Fig 4B). Cell cycle analysis revealed that IFNβ treatment was associated with a G0-G1 arrest (Fig 4E). Additionally, IFNβ was found to significantly reduce proliferation in both CR and SE cells, with the SE cells specifically increasing their doubling time from 17.5 hours to 22.5 hours (Fig 4F). These effects emerged progressively with continued cytokine exposure, consistent with stable cellular adaptation rather than a transient stress response. Thus, chronic and low-level IFNβ treatment phenocopies several aspects of chronic cisplatin treatment, similar to effects noted in our previous publication^22^.

**Figure 4:**
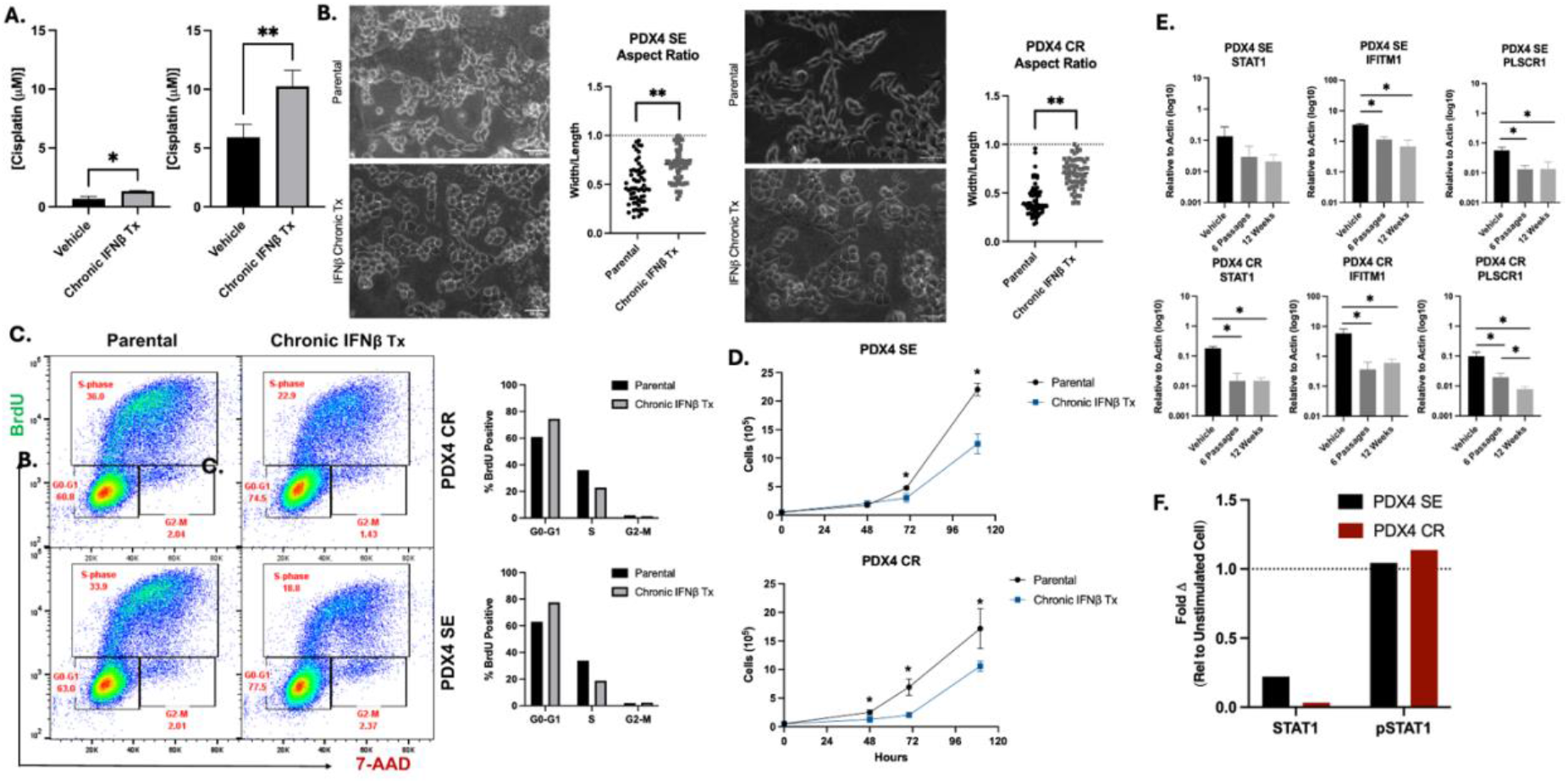
IFNβ treatment phenocopies chronic cisplatin treatment and drives therapy resistance. **A.** The IC50 of SE and CR cells was significantly higher following IFNβ treatment. **B**. Morphology change was determined by measuring the aspect ratio (width/legth) of 60 cells from each sample (20 cells per photo, 2 photos per well, 3 wells). Scale bar = 50µm. **C**. BrdU staining revealed cell cycle arrest in G0-G1 following IFNβ treatment, as measured by flow cytometry. Data represent one representative replicate. **D**. Proliferation was detected by counting individual wells of parental and IFNβ treated cells over time. **E**. RT-qPCR was used to analyze STAT1, IFITM1, and PLSCR1 following 6 passages and 12 weeks following consistent low-level IFNβ treatment. **F**. Intracellular flow cytometry was used to measure STAT1 and pSTAT1 protein levels following chronic IFNβ treatment. One representative replicate shown. Results are displayed as n=3, unless otherwise noted, and presented as the means ± SD. Statistical significance was determined using the unpaired t-test. p-values: p ≤ 0.05 (*) and p ≤ 0.01 (**).

Cells were harvested for analysis at both 6 passages and 12 weeks in order to monitor transcriptional changes associated with the phenotypic changes seen. Notably, chronic IFNβ exposure resulted in significant blunting of IRDS gene expression following both short-(6 passages) and long-term (12 weeks) IFN treatment (Fig 4E). On the protein level, STAT1 levels were notably decreased, while pSTAT1 levels were slightly increased (Fig 4F). These data indicate that the cells have exerted a negative regulatory mechanism, by which response to IFNβ has been suppressed.

In order to determine if the cells had been influenced into an anti-viral/anti-cancer state by the treatment, we measured the expression of CXCL10 (interferon-gamma-induced protein 10), a potent anti-viral/anti-cancer protein in ovarian cancer^40-42^ both following treatment with IFNβ, and following treatment with IFNβ and cisplatin, finding no increases among samples (Sup Fig 4A). We also evaluated the expression of several known regulatory proteins of the IFN-1 response pathway. IFNAR1 (interferon alpha receptor 1) expression mediates IFN-1 responses based on availability of the receptor for ligand binding and subsequent JAK/STAT activation^43^. USP18 (Ubiquitin-specific protease 18), is strongly induced by type 1 interferons and acts as a negative regulator of the IFN-1 response, showing that the roles of ISGs is dynamic, multifunctional, and tunable^44^. SOCS1 (suppressor of cytokine signaling 1) is a negative regulator is IFN-1 signaling, acting primarily to inhibit JAK/STAT signal transduction activation via interaction with IFNAR1 (interferon alpha receptor 1) associated Tyk2 (tyrosine kinase 2)^45^. IFNβ did not change the expression of any of these regulators, in CR or SE, indicating other mechanisms for ISG downregulation (Sup Fig 4A). We then considered epigenetic remodeling as a result of low-level IFNβ exposure. Epigenetic regulator KDM1B (Lysine-specific demethylase 1B) has been identified as an ISG that acts to mediate IFN-1 signaling^46^. This regulator was found to be significantly upregulated in SE, but not CR, cells following treatment with IFNβ, possibly suggesting a more complicated paradigm of IFN-1 response (Sup Fig 4B).

Together, these findings demonstrate that chronic, low-level IFNβ signaling is sufficient to recapitulate key features of cisplatin resistance in HGSOC cells, establishing IFNβ as a functional driver of resistance-associated phenotypes rather than solely a downstream

### IFNβ driven resistance is observed across additional HGSOC models

## DISCUSSION

Ovarian cancer represents a unique model of the development of therapy resistance. Most patients initially achieve partial or complete response, yet the majority ultimately recur. Recurrence following standard therapy is associated with therapy resistance and tumor aggression, indicating a causal link between treatment and recurrence^47,48^. DNA damage, from genetic instability and/or standard therapy, is associated with the enrichment of IFN-1 signaling and IRDS in HGSOC. However, whether IFN-1 plays a functional role in driving resistance and associated phenotypes or is merely a byproduct of chronic DNA damage remains unknown^49^. Our findings identify IFNβ as a functional driver of cisplatin resistance in HGSOC, capable of inducing phenotypic changes independent of the presence of a DNA-damaging agent and treatment-related IRDS upregulation.

IFN-1 as a marker of DNA damage, and even successful chemotherapy treatment, has been observed across multiple cancer types^17,39^. However, the precise role of IFN-1itself in driving resistance has not been established. The IRDS gene set has been heavily implicated in mediating DNA damage resistance and IFN-1 upregulation, with few specific mechanisms identified in ovarian cancer^17,18,20^. Our study aimed to clarify the role of IFN-1 in driving the phenotypes associated with therapy-induced DNA damage, while investigating if IRDS also plays a causal role. The stable IFN-1 activation and signaling phenomenon seen in PDX4 CR, compared to the SE counterpart, appears to be exclusive to cells chronically exposed to cisplatin over time. While RNA expression reflected this activation, the absence of corresponding robust protein secretion indicated low-level, cell-autonomous, autocrine-like signaling. We hypothesized that chemoresistant cells, with enrichment of these programs, would be “primed” to respond to the stress of therapy via robust IFN-1 signaling activation. However, upon acute treatment with cisplatin, an increase in STAT1, pSTAT1, and IFITM1 protein was observed in both SE and CR cells, indicating that a conserved IFN-1 response may exist. Of interest, however, the ratio of pSTAT1 to STAT1 expression in CR cells was greater than that in SE cells following treatment, indicating the “priming” response observed may only be relevant in terms of signaling downstream of IFN-1. Thus, a shift toward canonical signaling (ISGF3 instead of u-ISGF3) may play a role in phenotypic changes observed.

As shown, IFNβ treatment promoted the development of resistance while driving significant blunting of both canonical and non-canonical IFN-1 signaling programs, prompting further investigation into the precise mechanism of the promotion of therapy resistance. Most notably, STAT1 expression was significantly decreased in both CR and SE IFNβ-treated cells, suggesting its potential role in mediating resistance via altering signal transduction dynamics or availability. Separate and distinct negative regulators of IFN-1 responses were tested and revealed no changes in expression, indicating the blunting seen may be an epigenetic adaptation, as evidenced by increased expression of epigenetic regulator, KDM1B. Taken together, this evidence suggests that IFN-1 in the absence of DNA damage may contribute to resistance in an alternative, IRDS-independent manner.

The current paradigm of IFN-1 production in cancer suggests that DNA damage, driven by p53, or homologous recombination deficiency (HRD)-associated mutations, may be sufficient to drive chronic IFN-1 production and contribute to pro-tumor phenotypes, as suggested here^16^. In ovarian cancer specifically, localized IFN-1 and related signatures have been found to pre-exist^50^, being identified as early as emergence of p53 signatures, correlating with immune exhaustion as carcinoma progresses and increasing following chemotherapy exposure^19,51^. Consistent with this model, we identify IFN-1 and IRDS upregulation in CR cells, following chronic cisplatin exposure, as well as differential temporal upregulation of the cytokine in SE and CR cells following acute cisplatin exposure, with CR responding with latent, non-canonical signaling. These findings support the use of IFN-1 and related signaling genes as biomarkers of resistance risk and therapeutic response, serving a cautionary role for clinicians when selecting therapies for patients displaying an upregulation of these pathways.

Many pre-clinical and clinical studies observing and modulating the roles of IFN-1 production and signaling are focused on monitoring immune activation and overall tumor burden, rather than elucidating the precise roles of tumor cell-intrinsic IFN-1 signaling^52^. We focus here on systematically determining the role of IFNβ itself in promoting a pro-tumor phenotype. The limitations of this study include the use of a single patient-derived cell line (PDX4) in modeling developed therapy resistance and IFN-1 activation, and the in-vitro model itself. Future studies will incorporate the use of 3d systems and animal models to provide full context of the delicate dynamics of IFN-1 signaling, and will determine the factors that contribute to intrinsic IFN-1 activation, as well as determine specific mechanisms by which IFNβ itself promotes resistance.

In conclusion, we demonstrate that an IFNβ related signature is enriched in cisplatin-resistant HGSOC cells and is associated with latent IFN-1 cytokine production following acute cisplatin exposure. IFNβ was found to be a functional driver of therapy resistance, phenocopying other key features of chronic cisplatin exposure such as decreased proliferation, cell cycle arrest, and epithelial morphology changes. This evidence provides a basis for the continued research and development of biomarkers for therapy response and resistance, as well as further studies determining the underlying mechanism of IFNβ mediated resistance.

