## supplemental information for "Interferon Beta Drives Therapy Resistance in a Patient-Derived Model of High-Grade Serous Ovarian Cancer"

SUPPLEMENTAL DATA

Supplemental Methods

| **Primer** | **Forward (5'-3')** | **Reverse (5'-3')** |
| --- | --- | --- |
| Actin | TGAAGTGTGACGTGGACATC | GGAGGAGCAATGATCTTGAT |
| BST2 | ATGTCACCCATCTCCTGCAA | CGCGATTCTCACGCTTAAGAC |
| CXCL10 | CTGCCATTCTGATTTGCTGCC | AATGCTGATGCAGGTACAGCG |
| GBP4 | ATGGGTGAGAGAACTCTTCACG | TGCGGTATAGCCCTACAATGG |
| HERC6 | TTGCTGGAACATATGCCAAC | ACTTGCAGTCAGACAAGCAG |
| IFI6 | TGGTCTGCGATCCTGAATG | TACTTGTGGGTGGCGTAG |
| IFIT3 | CAGAACTGCAGGGAAACAGC | TGA ATAAGTTCCAGGTGAAATGGC |
| IFITM1 | CCCTGTTCAACACCCTCTTC | ATCCAATGGTCATGAGGATGCC |
| IFNAR1 | ATGTAACTGGTGGGATCTGCG | GTCGACCTCTACTTTTTGAGGAGA |
| IFNα (total) | GATGGCAACCAGTTCCAGAAG | CAGACAGGCTTCCAAGTCATTC |
| IFNα-1 | TTGACTCATACACCAGGTCACG | AGCATGGTCATAGTTATAGCAGGG |
| IFNα-2b | TGCAAGTCAAGCTTGCTCTGT | GGACAGGGATGGTTTCAGCC |
| IFNβ | CAATTGAATGGGAGGCTTGAATA | CAGTGCTAGATGAATCTTGTCTG |
| IFNε | GAGATGCTTCAGCAGATCTTC | TTCCAGGTAATCATGGATCCT |
| KDM1B | GCGTGATGTCTGTGATT | TTGTGGGATCTGGGACCT |
| MX1 | CTTTCCAGTCCAGCTCGGCA | AGCTGCTGGCCGTACGTCTG |
| OAS1 | TGAGGTCCAGGCTCCACGCT | GCAGGTCGGTGCACTCCTCG |
| PLSCR1 | AACTTGCCAGTTGGGTATCC | AGTTTAATGGAGGCTGTGGC |
| SOCS1 | TTTTCGCCCTTAGCGTGAAG | CATCCAGGTGAAAGCGGC |
| STAT1 | GACCCAATCCAGATGTCTATG | CCTTGTCCTTCACATTTCTGAC |
| USP18 | CAGAGGAGAAGCGTCCCTT | TCACCCGGATCGTATACAGG |

**Table 1.** RT-qPCR primers.

Supplemental Figures

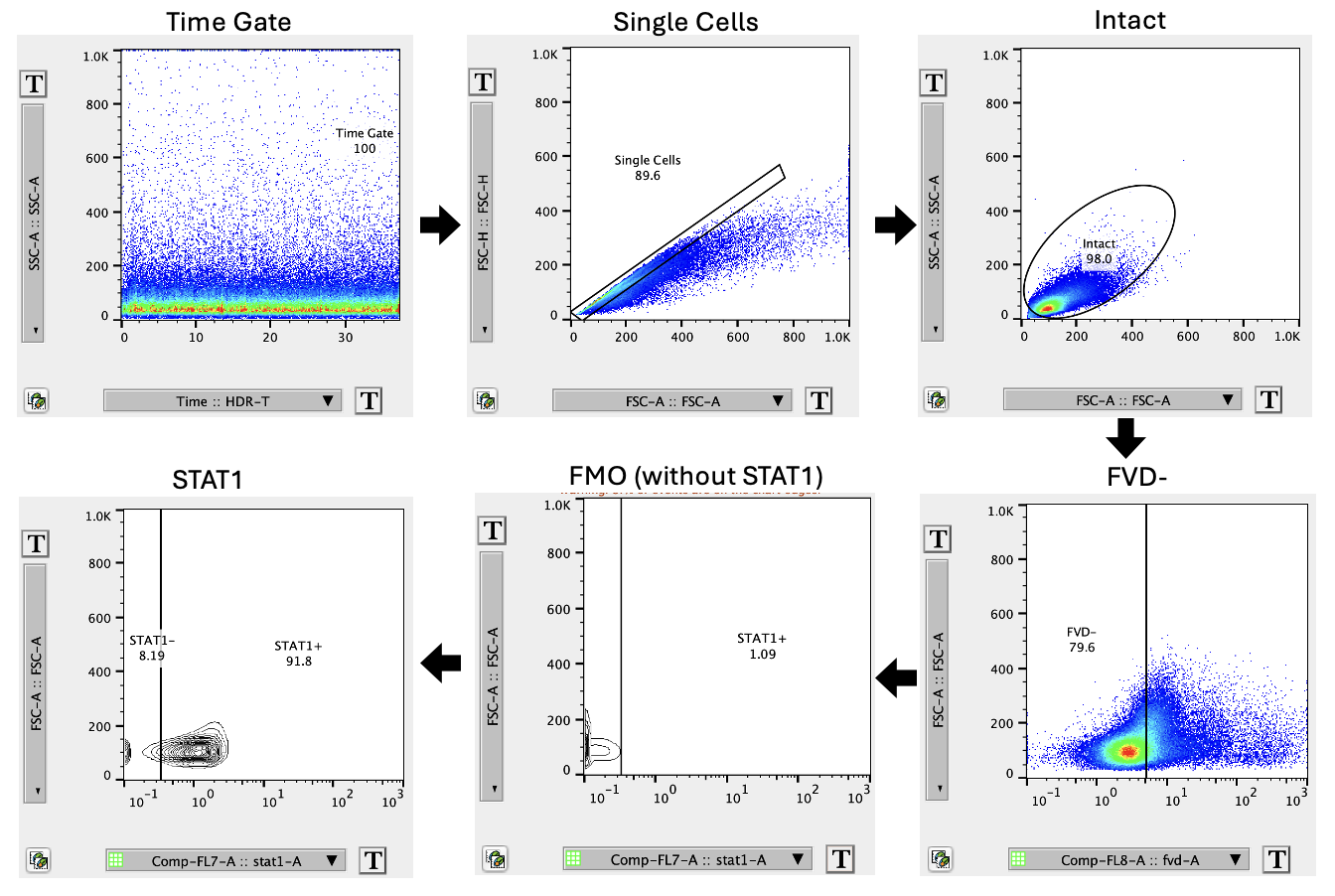

**Supplemental Figure 1**: Example gating scheme for flow cytometry. Time gate, single cell gating, intact gate, FVD negative gate, FMO negative and positive STAT1 gates, and STAT1 FMO gates applied to sample.

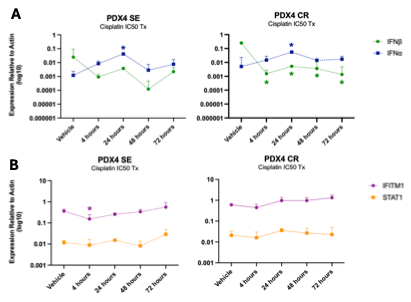

**Supplemental Figure 2.** **Temporal expression of IFN-1 and IRDS RNA following cisplatin treatment.** RT-qPCR of **(A.)** IFNα, IFNβ, **(B.)** STAT1, and IFITM1 in PDX4 SE and CR following 72 hours of cisplatin treatment at respective IC50 concentrations. Statistical analysis indicates comparison of listed time points to vehicle. Color of asterisk refers to the gene in reference for statistical comparison. Results are displayed as n=3, unless otherwise noted, and presented as the means ± SD. Statistical significance was determined using the unpaired t-test. p-values: p ≤ 0.05 (*).

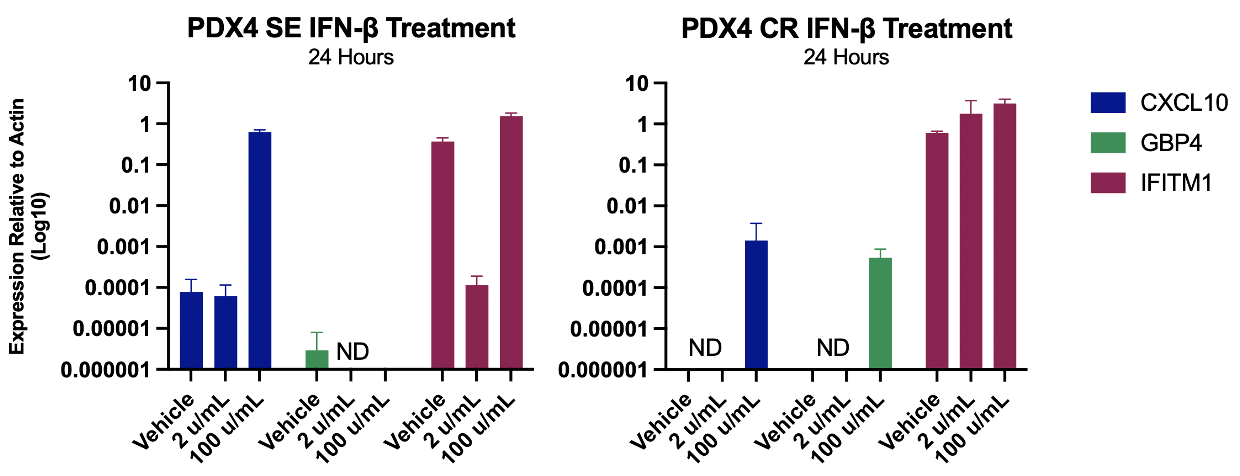

**Supplemental Figure 3. IFN**β **dose for generating acute responses.** Response of the cells to short-term (24 hours), low- or -high (2 or 100 U/mL) dose IFNβ was confirmed by measuring induction of the expression of an IRDS gene, IFITM1, as well as potent anti-viral/-cancer gene, CXCL10^1-3^, at both 2 U/mL and 100 U/mL. 2 U/mL did not induce immediate pro/anti-cancer effects, as measured by CXCL10 and IFITM1, while 100 U/mL promoted strong induction of CXCL10 and GBP4^4^ (CR only), indicating that 2 U/mL is appropriate for modeling chronic, low level IFN-1 siganling rather than acute immune-like activation.

ND = not detected. Results are displayed as n=3, unless otherwise noted, and presented as the means ± SD.

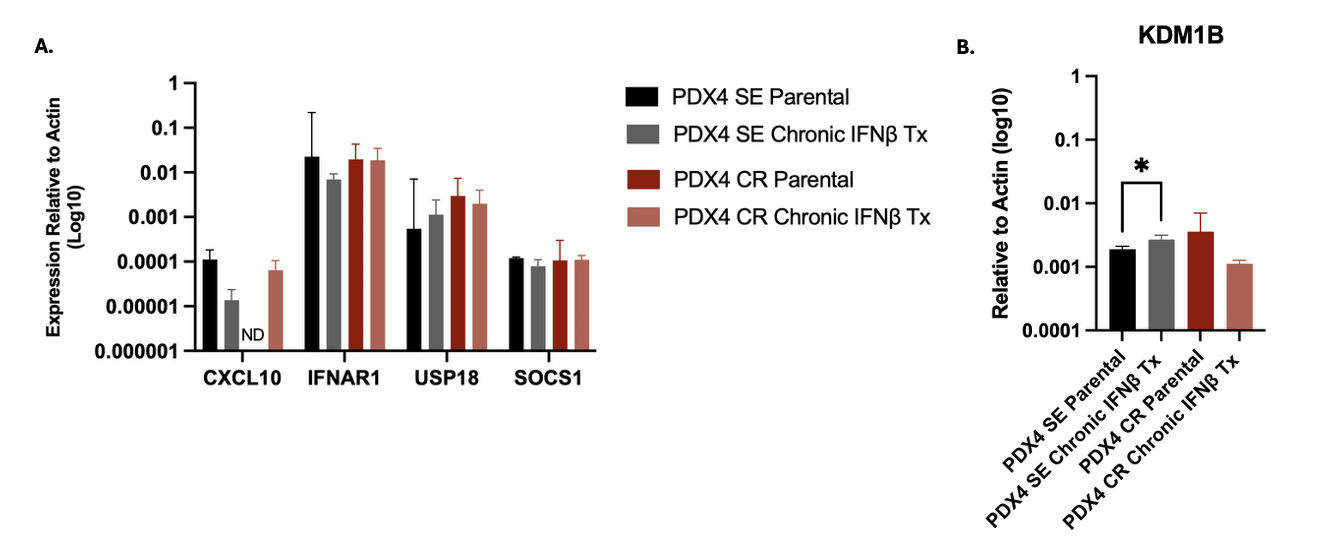

**Supplemental Figure 4: Validation of IFN-1 Signaling Regulatory Mechanisms. A.** CXCL10, IFNAR1, USP18, and SOC1 gene expression in PDX4 SE and CR **parental (darker) or chronic** IFNβ **treated cells (lighter) B.** KDM1B expression in PDX4 SE and CR **parental (darker) or chronic** IFNβ **treated cells. ND = not detetected.** Results are displayed as n=3, unless otherwise noted, and presented as the means ± SD. Statistical significance was determined using the paired t-test. p-values: p ≤ 0.05 (*).
